# Genes essential for embryonic stem cells are associated with neurodevelopmental disorders

**DOI:** 10.1101/567073

**Authors:** Shahar Shohat, Sagiv Shifman

**Affiliations:** Department of Genetics, The Institute of Life Sciences, The Hebrew University of Jerusalem, Jerusalem, Israel

## Abstract

Mouse embryonic stem cells (mESCs) are a key component in generating mouse models for human diseases and performing basic research on pluripotency, but how many genes are essential for mESCs is still unknown. We performed a genome-wide screen for essential genes in mESCs and compared it to human cells. We found that essential genes are enriched for basic cellular functions, are highly expressed in mESCs, and tend to lack paralog genes. Notably, we discovered that genes that are essential specifically in mESCs play a role in pathways associated with their pluripotent state. We show that 25% of human genes that are intolerant to loss-of-function mutations are essential in mouse or human ESCs and that the human phenotypes most significantly associated with essential genes are neurodevelopmental. Our results provide insights into essential genes in the mouse, the pathways which govern pluripotency, and suggest that many genes associated with neurodevelopmental disorders are essential at very early embryonic stages.

## Introduction

Essential genes are required for the organism survival or development. Recent advancement in CRISPR technology have allowed for the identification of essential genes in multiple human cancer cell lines (Tzelepis et al. 2016; Wang et al. 2017, 2015) and in human embryonic stem cells (hESCs) (Yilmaz et al. 2018). Human genes essential *in vivo* have been identified by searching for genes with lower than expected rate of functional mutations (Lek et al. 2016; Samocha et al. 2014). Surprisingly, the genes that are intolerant to mutations in the human population are associated with neurodevelopmental risk genes (Shohat et al. 2017; Samocha et al. 2014), but not with other extensively studied diseases, such as type 2 diabetes, early-onset myocardial infarction, inflammatory bowel disease, ulcerative colitis, or Crohn disease (Ganna et al. 2018).

Previous and ongoing efforts to detect essential genes in the mouse genome focused on generating and characterizing the phenotypes of knockout mice. These efforts generated knockouts for 4969 genes of which 1187 were found to be essential (Dickinson et al. 2016). However, these studies still do not cover the entire mouse genome, and for most of the essential genes they do not provide information regarding the developmental stage affected by the essential gene.

Here we report a genome wide screen for genes essential in mouse embryonic stem cells (mESCs), which we compared to similar screens in human cells and genes essential *in vivo.* Our analysis reveals the cellular pathways which are globally essential and those that are specific to mESCs. We found that genes essential specifically in mESCs affect pathways associated with their pluripotent state, and that neurodevelopmental disorders are the human phenotypes most significantly associated with essential genes.

## Results

### CRISPR screen in mouse embryonic stem cells

We performed a loss-of-function genome-wide screen to detect genes essential for the survival and proliferation of mESCs. We used a pooled knockout CRISPR library that consists of 77,637 guide RNAs (gRNAs) targeting 19,674 coding genes (4 gRNA per gene) and 1000 non-gene targeting control gRNA (Doench et al. 2016). Sequencing of the pre-transfected plasmid library revealed the presence of 99.82% of gRNAs (143 gRNAs were missing) and that all genes had at least 97 reads from gene-targeting gRNAs (a mean of 1087 reads per gene) (Figure S1 A&B, Table S1). We transfected Cas9 expressing mESCs with lentiviruses containing the library and allowed the cells to proliferate for 18 days. Cells were collected on days 8, 11, 15 and 18 post transfection and the abundance of gRNAs was determined by sequencing (Figure 1A). In order to determine which genes are under significant negative selection we developed an approach that is based on a simulation of randomly selected control gRNAs (see Methods). This approach allows detection of negative and positive selection in the presence of random changes in gRNAs abundance (random drift) (Figure S1 C&D). Using this method, we were able to detect 2379 negatively selected genes, which are essential for survival or proliferation of mESCs (Figure 1B, Table S2).

**Figure 1.**
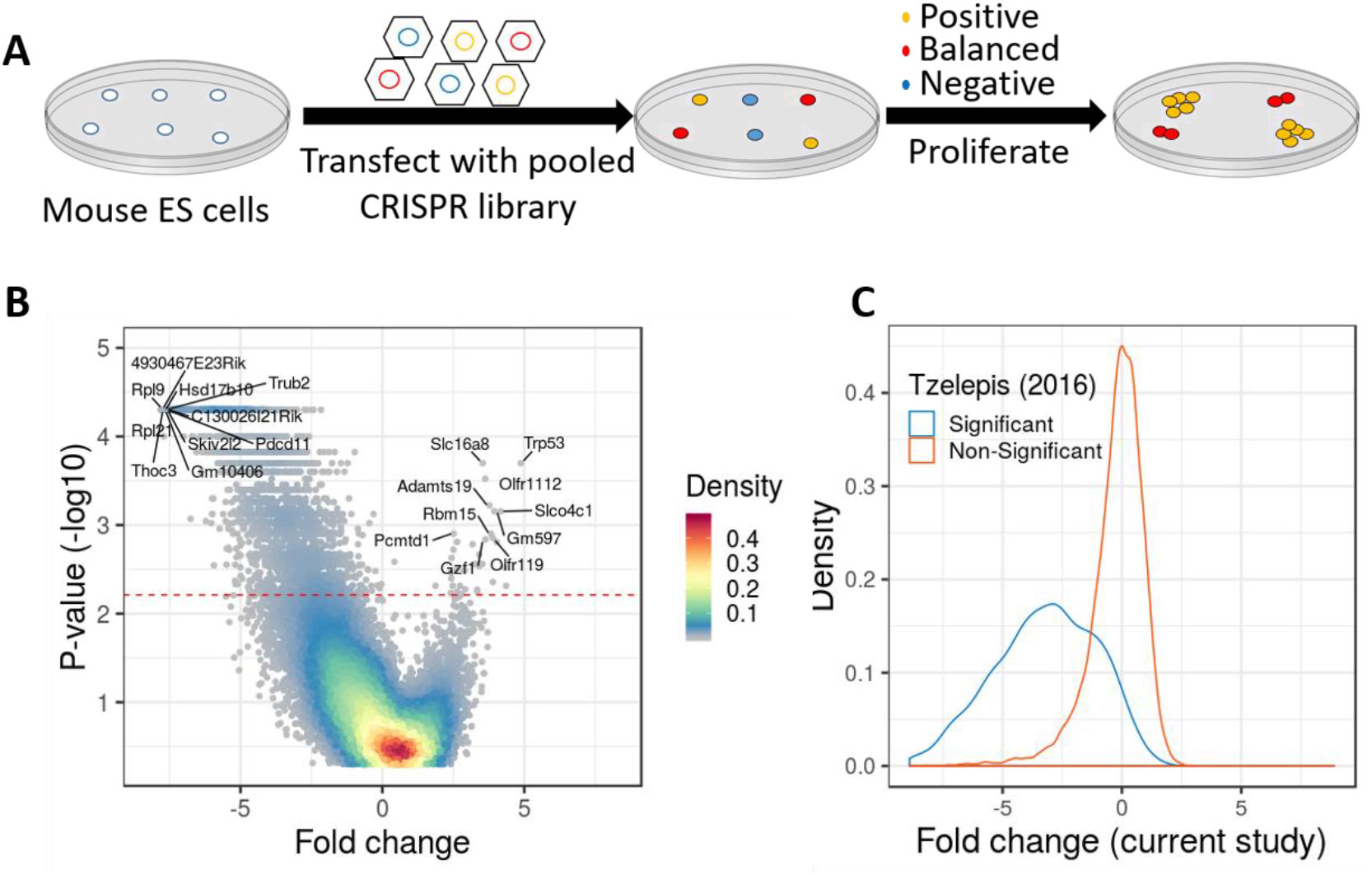
A Genome-wide CRISPR screen to identify genes essential in mESCs. **(A)** Schematic overview of the genome-wide CRISPR screen. **(B)** A volcano plot of the screen results. Significance (-log10 of the p-value) across all days as a function of the average fold change for each gene at the end of the screen. Color indicates the density of the points. The red line represents a corrected *P* < 0.05. **(C)** The distribution of the average fold change per gene (day 15) was drawn separately for genes with significant evidence for negative selection in a previous study (Tzelepis et al. 2016) (blue) and genes with no significant evidence for selection (orange).

Since results from a single screen might be influenced by the specific CRISPR library used or by other factors, we compared the results to data from another mESC line transfected with a different CRISPR library (Tzelepis et al. 2016). Despite the different experimental conditions, we found a highly significant overlap (odds ratio [OR] = 27, *P* < 10^−16^). Quantitatively, we observed that the distribution of fold-changes for gRNAs negatively selected in one screen was significantly shifted in the second screen, towards the same direction (*P* < 10^−16^) (Figure 1C). Since the two screens showed a very significant overlap, we combined the evidence of the two independent screens and generated a consensus list of genes under selection in mESCs, with 2164 genes under negative selection (henceforth, mESCs essential genes) (Table S3).

### mESCs essential genes are enriched with fundamental cellular processes

We next wanted to characterize the list of essential genes and test which biological process are involved. Using gene ontology (GO) and KEGG pathways enrichment analysis we found that the essential genes are associated with fundamental cellular processes such as ribosome biogenesis, RNA processing, translation, DNA metabolism and cell cycle (Figure 2A, Figure S2A & Table S4). For some KEGG pathways (Ribosome, DNA replication and RNA polymerase) nearly all genes (>90%) in the pathway were found to be essential (Figure 2A). The fact that not all genes in the top enriched KEGG pathways (“essential pathways”) immerged as essential in our screen could be a result of statistical power but another reason could be that some genes within the pathways are robust to mutations. One option for this robustness is functional redundancy, when an alternative gene or pathway can compensate for the mutated gene. The most likely candidates for this compensation are paralog genes. To investigate this possibility, we tested within each essential pathway if genes with paralog are less essential. For 7 out of the 10 essential pathways, genes with paralogs were significantly less essential compared to genes without paralogs (Figure 2B).

**Figure 2.**
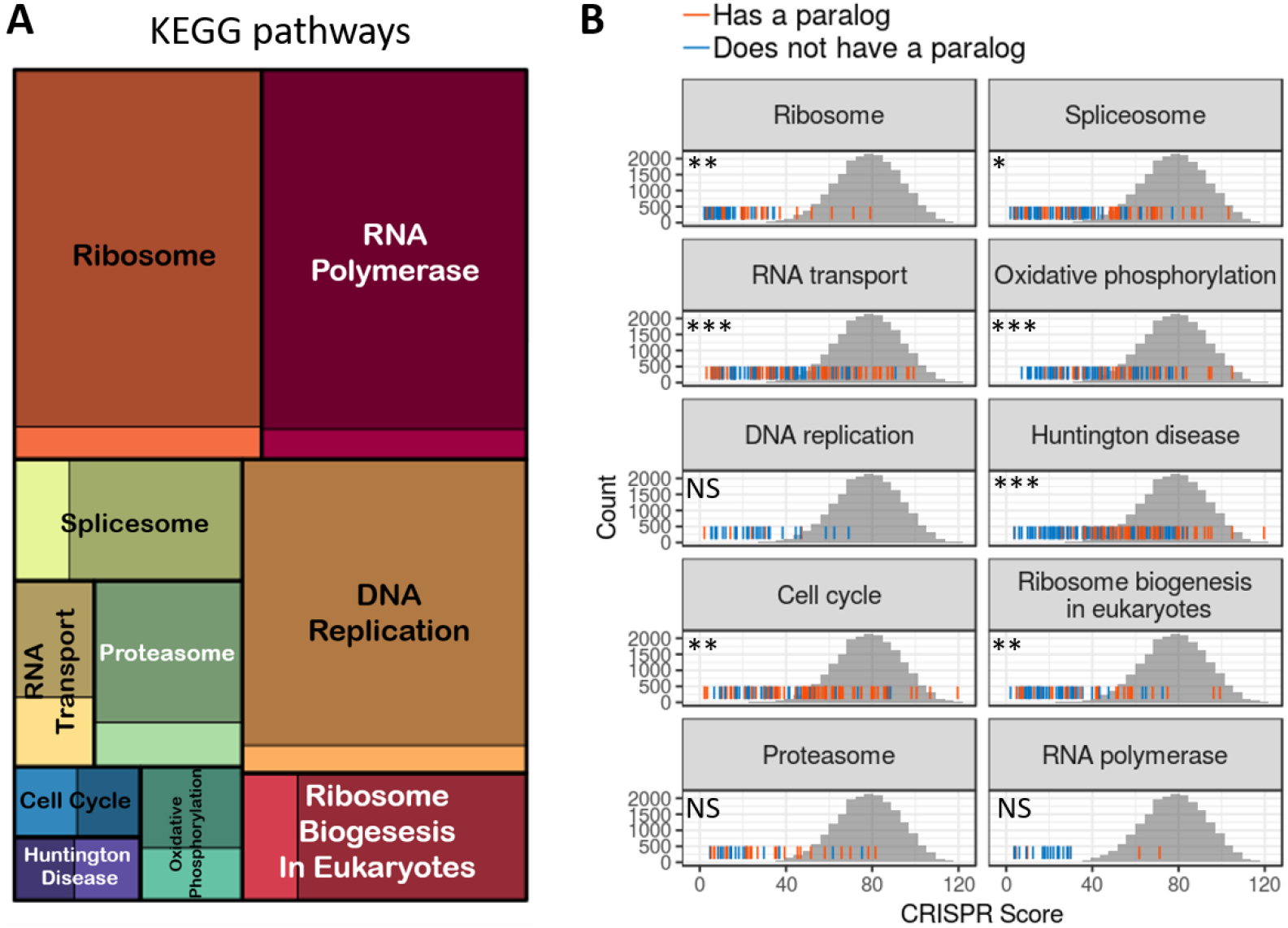
Essential genes belong to basic cellular pathways and tend to lack paralogs. **(A)** A treemap of the top 10 most enriched KEGG pathways (essential pathways). The square size is proportional to the enrichment strength and the color intensity indicates the proportion of essential (dark color) and non-essential genes (bright colors) in the pathway. **(B)** For each essential pathway the rug plot displays the distribution of CRISPR scores (sum of ranks across days) for all genes in the pathway. Orange lines indicate genes with a paralog, and blue lines indicate genes without a paralog. The grey histogram behind is the empirical null distribution of CRISPR scores based on control gRNAs. Significance indicates the differences in scores between genes with and without a paralog. NS, non-significant (*P* > 0.05); *, *P* < 0.05; ** *P* < 0.01; *** *P* < 0.001.

We found 115 genes (5% of the essential genes) in our screen that are essential but currently not associated with any GO term or pathway. To exclude the possibility that those genes are false positives we tested their expression in mESCs. We found that essential genes lacking a GO term, show significantly higher expression in mESCs than non-essential genes (*P* <10^−16^, Figure S2B). This indicates that there is a large list of uncharacterized genes with unknown functions that are essential in mESCs and probably essential for the survival of early-stage embryos.

### Essential genes with faster gRNA depletion have higher expression in mESCs and a lower number of paralogs

Our screen includes multiple time points allowing us to quantify the kinetics of the negative selection. When fold change across time points was used in hierarchical clustering, most genes were mapped into two main clusters: a fast-declining cluster (55%) that had a rapid reduction in gRNA representation in the first eight days (Figure 3A & S3A), and a gradual-declining group of genes (17%) with a linear decline of gRNAs across all days (Figure 3B). To investigate possible reasons for the two clusters we first tested if the gradual-declining group code for more stable proteins. Since data on protein stability in mESCs was unavailable, we used data from six other human or mouse cell lines as a proxy. We observed no significant difference in protein half-life between the groups (Figure S3B). Another, possibility is that the gradual-declining group are less essential. In agreement, we found that genes in the gradual-declining group showed significantly lower expression in mESCs (Figure 3C, *P* = 10^−5^) and significantly more paralogs than those in the fast-declining group (*P* = 0.027 Figure 3D). When examining the enrichment of cellular processes in the two gene groups, we found that the fast-declining genes showed a stronger enrichment for ribosome biogenesis, transcription and RNA processing terms. In contrast, the gradual-declining genes showed a higher enrichment for mitochondrial terms, and terms related to modifications of DNA and proteins (Figure S3 C-F). These results suggest that the difference in the selection rates mostly reflects differences in the essentiality level.

**Figure 3.**
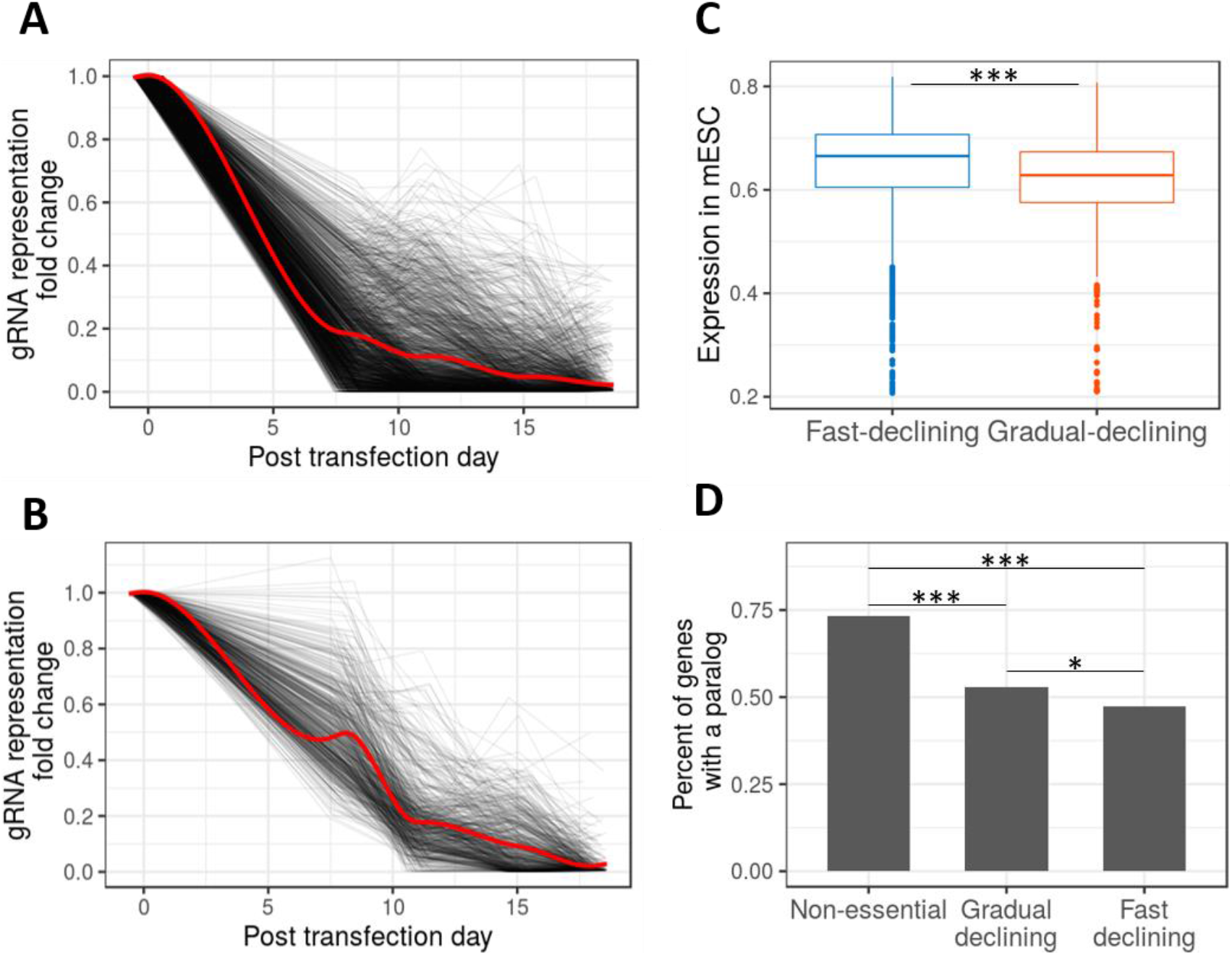
Gene essentiality as a quantitative phenotype. **(A, B)** Two different dynamics of gRNA depletion rates (see Figure S3A for clustering results): **(A)** fast-declining cluster, and **(B)** gradual-declining cluster. For each gene the gRNA with the strongest decline is shown (black lines). The red line indicates the trend of all genes in the cluster. **(C)** Expression in mESCs of genes in the fast and gradual declining groups. Expression values are log10 probe signal intensities from microarray data. **(D)** Percentage of genes with at least one paralog gene. *, *P* < 0.05; ***, *P* < 0.001.

### Genes essential specifically in mESCs are associated with the early pluripotent state

Essential genes in mESCs are enriched for the most basic cellular functions, similar to findings in other cell types (Yilmaz et al. 2018; Wang et al. 2017). We were interested to discover if there are genes that are essential specifically in mESCs. To this end, we compared the list of essential genes in mESCs to human cancer cell lines (n=11) and human haploid ESCs (n=1). We found that most genes negatively selected in mESCs are also essential in the cancer cell lines: 83% of the essential genes in mESCs were also identified in at least one out of the eleven human cancer cell lines and 39% were common to six or more cell lines (Figure 4A). When compared to hESCs, we found that 52% of the mESCs essential genes are also essential in hESC. To further assess the degree of agreement between mESCs and the human cells, we ranked the genes in each screen based on the negative selection level and compared the distribution of ranks between mESCs vs. all other screens. This analysis, which accounts for differences in statistical power between screens, further showed a highly significant agreement between mESC and human cells (Figure S4A).

**Figure 4.**
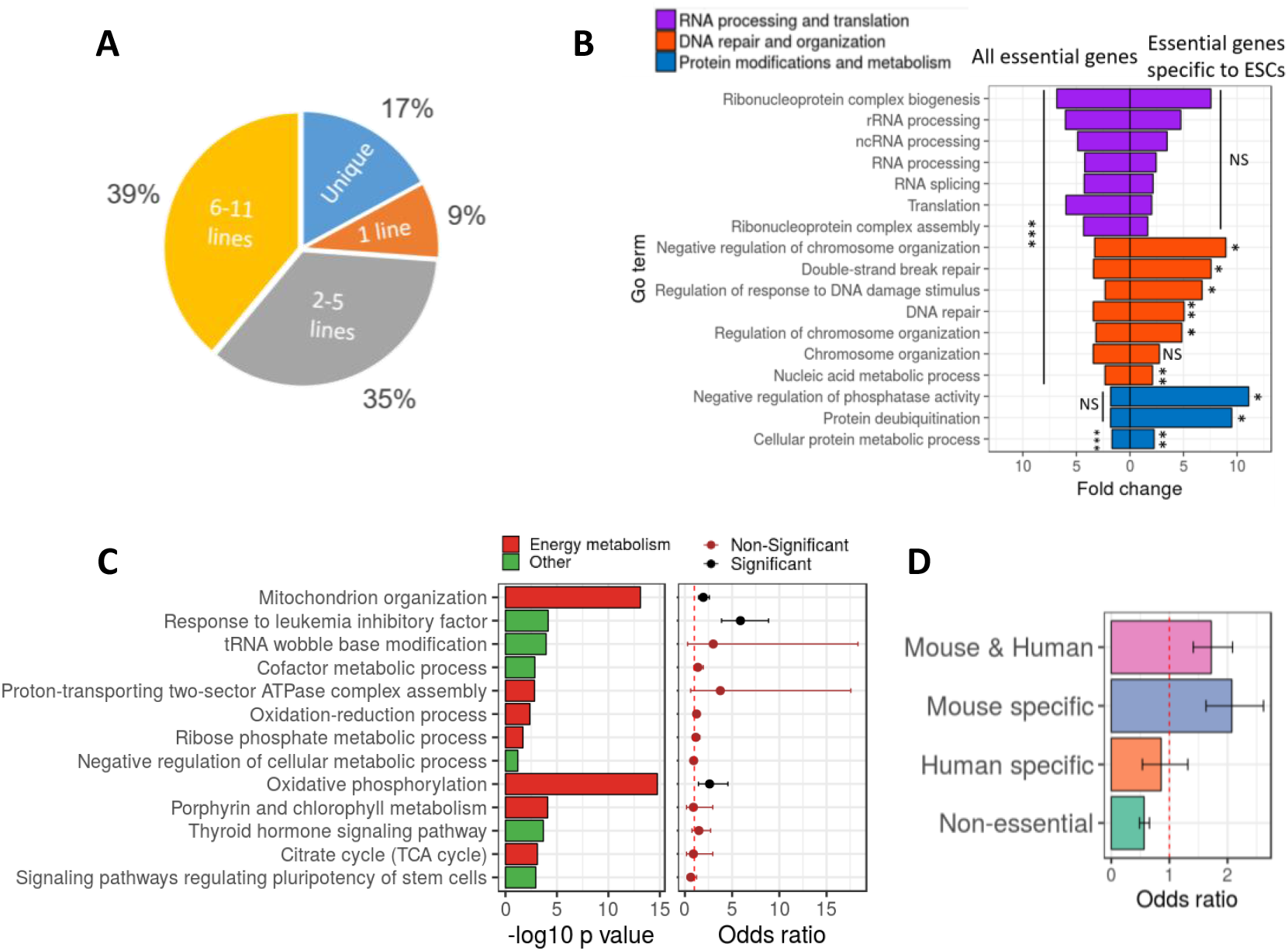
Analysis of genes essential specifically in mESCs reveals pathways associated with the pluripotent state. **(A)** The percentage of genes essential in mESCs which are also essential in one or more human cancer cell lines. **(B)** Comparison between GO term enrichment analysis for all genes essential in mESCs (left side) and genes specifically essential in human and mouse ESCs (right side). **(C)** Significant enrichment of GO terms and KEGG pathways for genes essential specifically in mESCs (left plot), and corresponding association level of those terms with genes significantly upregulated in mESCs relative to EpiSCs (right plot). Values are odds ratio ± 95% confidence interval. **(D)** Association analysis between genes upregulated in mESCs and (1) genes essential specifically in both human and mouse ESCs, (2) genes essential specifically in mESCs, (3) genes essential specifically in hESCs, and (4) non-essential genes. Values are odds ratio ± 95% confidence interval. NS, non-significant (*P* > 0.05); *, *P* < 0.05; ** *P* < 0.01; *** *P* < 0.001.

Although most essential genes are identified also in other cell types, some are specific to ESCs (both human and mouse) and some are essential only in mESCs. Genes essential specifically in ESCs were enriched for GO terms related to DNA repair and organization, and for protein modification, but were not significantly enriched for RNA processing and translation, which were the top enriched terms when considering all essential genes (Figure 4B). Genes essential specifically in mESCs showed an enrichment for mitochondria and other energy metabolism terms (Figure 4C right). The strongest enrichment was in the oxidative phosphorylation pathway, where complexes 1,2 and 4 were extremely specific to mESCs essential genes, and complexes 3 and 5 contain genes essential in both mouse and human ESCs (Figure SSB). In addition, cellular response to leukemia inhibitor factor (LIF) and signaling pathways regulating pluripotency of stem cells were also specifically enriched with mESCs essential genes (Figure 4C right). Genes essential specifically in hESCs showed enrichment in the autophagy pathway, cholesterol biosynthesis and in the terpenoid backbone biosynthesis (Figures S5A). It is interesting to note that the enrichment in the terpenoid backbone biosynthesis pathway is concentrated in the Mevalonate pathway (which is also related to cholesterol synthesis), and that all the key enzymes in the Mevalonate pathway are essential only in hESCs (Figures S5B).

The enrichment for LIF signaling, and energy metabolism related terms implies that genes essential specifically in mESCs might be related to the pluripotent state of mESCs and not to the difference between mouse and human. To test this hypothesis, we identified genes differential expressed (DE) between mESCs (early pluripotent) and mouse epiblast stem cells (EpiSCs) (represent primed pluripotent cells) (Brons et al. 2007; Tesar et al. 2007) and tested the enrichment of the DE genes in pathways/terms associated with mESCs-specific essential genes. Out of the 13 KEGG and GO terms enriched with mESCs essential genes, 3 were significantly enriched for genes upregulated in the early pluripotent state (response to LIF, mitochondria organization and oxidative phosphorylation) (Figure 4C left). When considering all essential genes, we found that genes essential in both human and mouse ESCs, and genes essential specifically in mESCs are significantly associated with genes upregulated in the early pluripotent state (*P* = 1.1×10^−7^, OR = 1.7; *P* = 5×10^−9^, OR = 2; respectively). Genes essential specifically in hESCs showed no significant association with genes upregulated in the early pluripotent state (*P* = 0.53, OR = 0.85), and non-essential genes were significantly under enriched (*P* = 2.8×10^−13^, OR = 0.56) (Figure 4D).

### Essential genes in mESCs are intolerant to heterozygote and homozygote mutations

Our screen is based on mESCs proliferating *in vitro* and thus the essential genes may not all be required *in vivo.* To test if the genes identified in the screen are known to be essential *in vivo,* we first compared the list with genes previously associated with embryonic lethality in mice. We found that 71% of the essential genes in mESCs (with phenotypic information) are known to be embryonic lethal (OR = 13.05, *P* < 10^−16^), and additional 21% are known to be postnatal lethal or cause abnormal growth (Figure 5A). Based on preferential gene expression, the 8% of genes without developmental phenotypes are likely to be true essentials. Since the genes without developmental phenotype show significantly higher expression in mESCs when compared to non-essential genes (*P* < 10^−16^), and are non-significantly different from essential genes with developmental phenotypes (*P* = 0.17) (Figure S6A). Next, we tested the overlap of genes essential in mESCs with genes intolerant to loss-of-function (LoF) mutations in humans (LoF intolerant) (Lek et al. 2016). We found that 30% of the essential genes in mESCs are LoF intolerant in humans (OR = 1.9, *P* < 10^−16^) (Figure 5B). The relatively low overlap with LoF intolerant genes may be due to the differences between mouse and human, however, the percentage of overlap between hESCs essential genes and LoF intolerant genes was only slightly higher (30% vs. 35%; *P* =0.01) (Figure 5B). Given the similar level of magnitude in overlap with LoF intolerant genes, we combined the two list of essential genes in mESCs and hESCs to one list of ESCs essential genes.

**Figure 5.**
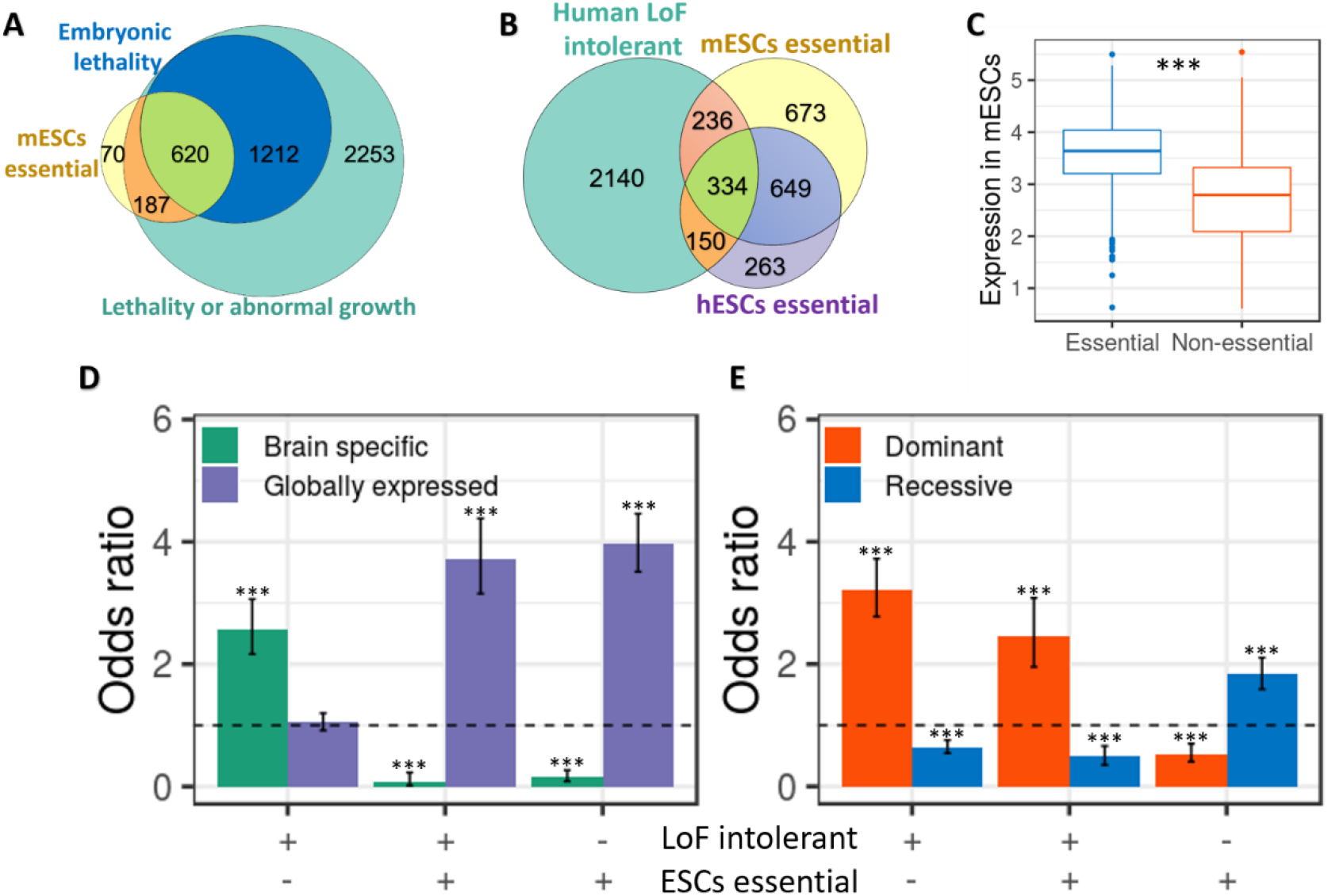
Overlap between essential genes in mESCs and genes essential *in vivo* in mouse and human. **(A)** Overlap between essential genes in mESCs and embryonic lethal genes in mice (odds ratio = 12.84, *P* < 10^−16^), and with genes known to be lethal or cause abnormal growth (odds ratio = 13.94, *P* < 10^−16^). **(B)** Overlap between human LoF intolerant genes and essential genes in mESCs (odds ratio = 1.88, *P* < 10^−16^) and essential genes in hESCs (odds ratio = 2.33, *P* < 10^−16^). **(C)** Differences in expression level in mESCs for human LoF intolerant genes divided to genes that are essential or non-essential in ESCs. Expression values are signal intensities (log10) from microarray data. **(D)** Association between the pattern of expression (brain specific genes vs. globally expressed genes) and being essentiality in ESCs and/or intolerant to LoF in humans. Values are odds ratio ± 95% confidence interval. **(E)** Association between dominant and recessive inheritance and essentiality in ESCs and/or intolerant to LoF in humans. Values are odds ratio ± 95% confidence interval. *** *P* < 0.001.

One possible explanation why many LoF intolerant genes are not essential in ESCs is that they are essential in later developmental periods. In accordance, we found that LoF intolerant genes that are not essential in ESCs express at relatively low levels in mESCs (*P* <10^−16^, Figure 5C). In addition, when testing which tissues are most influenced by these genes, we found that LoF intolerant genes that are not essential in ESCs, preferentially express in the brain, while genes that are both LoF intolerant and essential in ESCs are broadly expressed across multiple tissues (Figure 5D). The relatively low overlap between LoF intolerant genes and genes essential in ESCs also raises the question of why many genes that are essential in ESCs are not LoF intolerant. We hypothesized that LoF intolerant genes represent genes sensitive to heterozygote mutations, while in CRISPR screens LoF mutations are often formed in both copies of the gene. Indeed, we found that the essential genes that are also LoF intolerant are more likely to be associated with dominant disorders, while essential genes that are tolerant to LoF mutations are more likely to be associated with recessive disorders (Figure 5E).

### Essential genes in ESCs are associated with neurodevelopmental phenotypes

Since ESCs serve as a model for studying many human diseases (Keane et al. 2014; Zhu and Huangfu 2013), we tested what type of human phenotypes associated with genes essential in ESCs. When testing all human phenotypes in the Human Phenotype Ontology (HPO) (Köhler et al. 2019), we found that essential genes are enriched for abnormal phenotypes related to the brain, muscle or blood systems (Figure 6A). Notably, there is a large overlap between the list of genes belonging to different phenotypes, where most genes belong to more than one phenotype, and 92% are associated with brain related phenotype (Figure 6SB). Next, we tested if essential genes associated with human diseases are enriched with GO term that are different than the general GO term of essential genes. We found that essential genes associated with diseases are significantly more enriched for energy related GO terms, for aminoacyl-tRNA biosynthesis, and are significantly less enriched for the proteasome (Figure 6SC).

**Figure 6.**
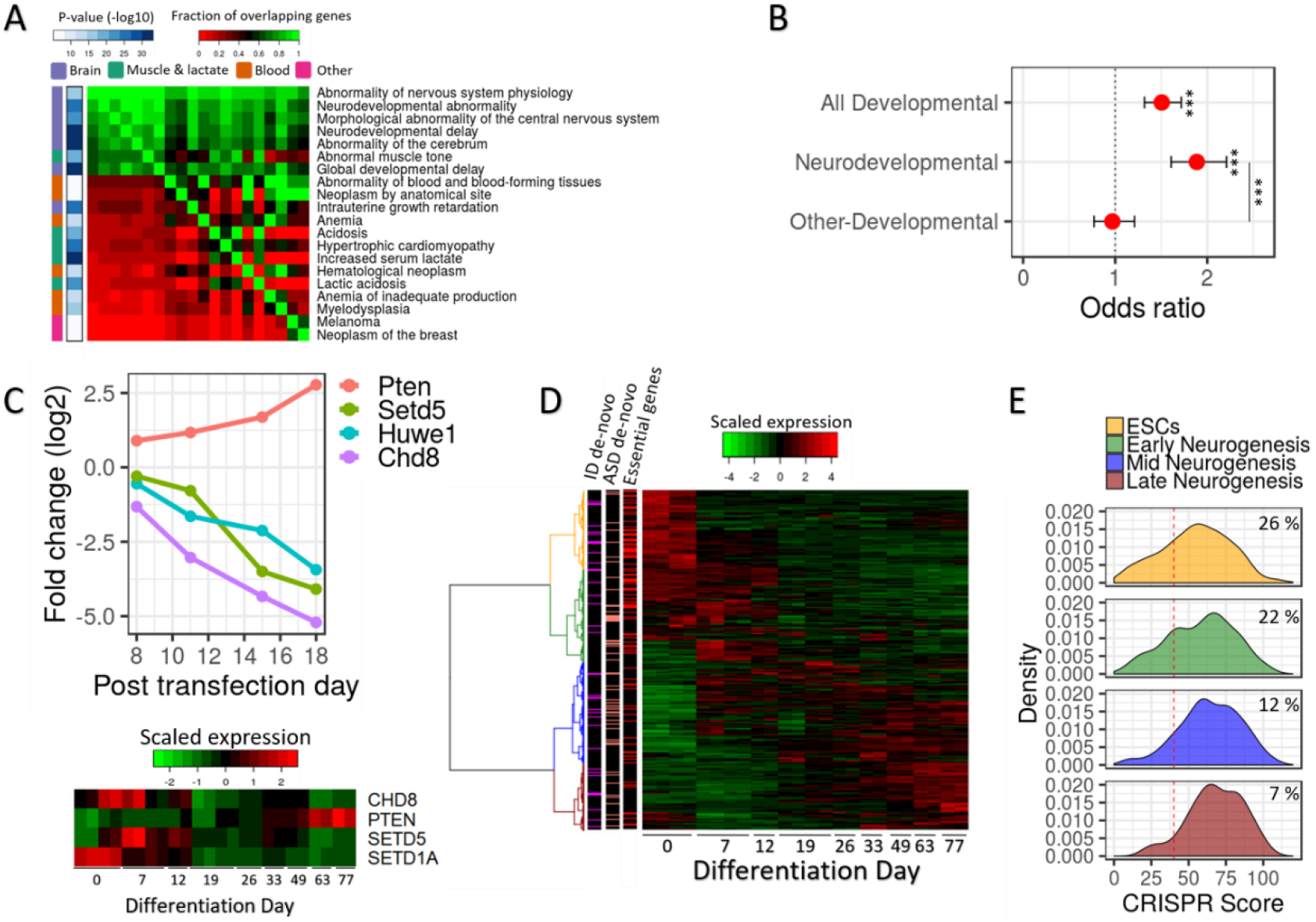
Essential genes in ESCs are associated with neurodevelopmental phenotypes. **(A)** Human phenotypes significantly enriched with essential genes. The phenotypes are ordered by the degree of overlap with other phenotypes. The color in the heatmap corresponds to the degree of overlap in genes between phenotypes (red is low overlap; green is high overlap). The right-side bar indicates the significance level of the enrichment of each phenotype with essential genes. The left side bar indicates the organs involved in the phenotypes. **(B)** Association between ESC essential genes and human developmental and neurodevelopmental disorders. Values are odds ratio ± 95% confidence interval. ***, corrected *P* < 0.001. **(C)** (Top) Dynamic of gRNAs (average fold change) targeting four genes associated with neurodevelopmental disorders. (Bottom) The expression of the four genes during human *in vitro* corticogenesis. The color of the heatmap correspond to the normalized expression (red is high and green is low levels of expression). **(D)** A heatmap of gene expression of risk genes for neurodevelopmental disorders during human *in vitro* corticogenesis. The color of the heatmap correspond to the normalized expression (red is high and green is low levels of expression). Side bars indicate ESCs essential genes (red), genes implicated in ASD by multiple *de novo* mutations (salmon) and genes implicated in ID by multiple *de novo* mutations (magenta). **(E)** Distribution of CRISPR scores for NDD risk genes divided by expression patterns during human *in vitro* corticogenesis. The red line indicates a corrected *P* < 0.05. The numbers are the percentage of ESCs essential genes in each group.

The association between genes essential in ESCs and human phenotypes may be related to the role of those genes in regulating proliferation and differentiation during development, therefore we studied the association between genes essential in ESCs and human developmental disorders, including neurodevelopmental disorders (NDDs). We tested the overlap with genes previously identified to be disrupted by *de novo* mutations in individuals with different developmental phenotypes (Wright et al. 2015). Out of the list of essential genes in ESCs, 14% were previously identified to be mutated in individuals with developmental phenotype (OR = 1.5, *P* = 2.5×10^−9^, Figure 6B). We separated the individuals to ones with neurodevelopmental phenotypes and ones without brain-related phenotypes, and found that the genetic overlap with ESCs essential genes was driven only by the neurodevelopmental phenotypes (OR_NDDs_ = 1.88, *P* _NDDs_ =2.6×10^−14^; OR non-NDDs = 0.97, *P* _non-NDDs_ = 0.8; difference between NDDs and non-NDDs, *P* = 2.9×10^−6^). Among the NDD risk genes that were identified as ESCs essentials are well-known genes for autism spectrum disorder (ASD) (CHD8) (Bernier et al. 2014), schizophrenia (SETD1A) (Bernier et al. 2014) and intellectual disability (ID) (SETD5) (Bernier et al. 2014) (Figure 6C).

Our results suggest that many genes associated with NDDs are essential as early as the ESC stage. We predicted that those NDD genes that are essential in ESCs will be preferentially expressed in ESCs, since in general, genes essential in ESCs were preferentially expressed in human and mouse ESCs (Figure S6D). To study NDDs genes, we tested the expression patterns of NDDs risk genes and specifically ASD and ID risk genes during human and mouse *in vitro* corticogenesis. While ASD and ID risk genes are preferentially expressed across multiple differentiation stages (Figure 6D, Figure S6E), those genes that show the highest expression, in both human and mouse, at the embryonic stem cell stages have a significant propensity to be essential in ESCs (Figure 7D, Figure S6F).

## Discussion

CRISPR screens have been used to identify essential genes mainly in human cancer cells but less commonly in other types of cells and organisms. We have shown here that a screen in mESCs can identify not only mouse essential genes and genes involved in pluripotency, but also genes related to human disorders. Our results raise issues about the quantitative definition of essential genes, and about the biological insights that can be attained by comparing essential genes identified *in vivo* and *in vitro.*

Our analysis shows that essential genes, are less likely to have a paralog gene. This phenomenon was observed in yeast (Keane et al. 2014) and C. elegans (Kamath et al. 2003), but was debated in the mouse (Liao and Zhang 2007; Makino et al. 2009; White et al. 2013). This debate was based on limited number of genes and biased screens. Our analysis, which covers the vast majority of mouse genes, confirms that essential genes are less likely to have a paralog across the eukaryotic tree.

The screen we performed spans over several time points, which allowed us to quantify the gRNA decline rate and cluster the essential genes into two groups. We found that genes in the fast declining group are more highly expressed, less likely to have a paralog and are enriched for fundamental cellular functions, all of which are general properties of essential genes (Rancati et al. 2017; White et al. 2013). While there is possibility that the genes in the fast-declining group are essential for survival whereas the genes in the gradual-declining group are essential for proliferation, our analysis implies that there is a quantitative difference in the level of essentiality between the gene groups.

Our results imply that the differences between essential genes in human and mouse ESCs are mostly related to differences in their pluripotent state and not to the organism. We show that genes which are specifically essential in mESCs are included in pathways associated with the pluripotent state, such as LIF, mitochondria organization, and oxidative phosphorylation (Dahéron et al. 2004; Zhou et al. 2012; Brons et al. 2007), while genes essential specifically in hESCs are involved in cholesterol synthesis. Cholesterol has been previously shown to play a role in cell cycle regulation (Singh et al. 2013; Wang et al. 2018), and a recent study found that cholesterol synthesis is upregulated in primed pluripotent cells when compared to naïve cells (Sperber et al. 2015). Our analysis show that cholesterol synthesis is not only upregulated in primed ESCs but is specifically essential to cells in this state of pluripotency. Since the screen in hESCs was performed with haploid cells, it is possible that some of the differences we observed might in part be connected to the haploid state. Ideally, a direct comparison between diploid cells from the same specie would be required to resolve the complex regulation of different pluripotency stages.

Our study provides an important new resource for studying human disease. Genes known to be intolerant to LoF mutations in humans are mainly dominant acting essential genes. For most of those LoF intolerant genes it is still unknown which developmental period is affected. Our screen provides a list of essential genes that includes recessive and dominant inheritance, and shows that 25% of human LoF intolerant genes are essential already at the ESCs stage. The remaining 75% show lower expression in ESCs and are likely essential in later stages of development. When testing which human disorders are associated with essential genes in ESCs we found that brain developmental disorders are the most significantly enriched. Since human genes intolerant to LoF mutations are also associated with neurodevelopmental disorders (Shohat et al. 2017; Samocha et al. 2014), it raises the question why essential genes are not linked to diseases affecting other important organs. We suggest that this could be related to the inheritance pattern, since mutations disrupting both copies of essential genes will frequently lead to embryonic lethality and therefore will not be linked to any human disease. However, it is possible that during development the brain is the most sensitive organ to heterozygote LoF mutations in essential genes, leading to deficits in proliferation and differentiation (Ernst 2016; Stephenson et al. 2011; Courchesne et al. 2003).

In summary, our work provides a map of genes essential for mESCs proliferation and survival. This data greatly expands our knowledge about genes essential in the mouse, genes essential for pluripotency, and can help researchers determine how and which human disorders can be modeled in the mouse. It is noteworthy in this regard that 5% of the essential genes are without any known function, and many others have very limited functional information. These genes are likely to be of great interest for further study.

## Methods

### Cell Culture

mESCs were a provided by A. Smith (Ying et al. 2003). Cell were cultured at 37 °C and with 5% CO2 on gelatin coated plates in mESC growth medium composed of knockout Dulbecco’s modified Eagle’s medium (DMEM) supplemented with 15% ES cell-qualified Fetal bovine serum, 0.1 mM non-essential amino acids, 100 μM β-mercaptoethanol, 50 μg ml-1 penicillin, 50 μg ml-1 streptomycin, 100unitsml-1 Leukemia inhibitory factor, 2 mM L-glutamine, 3uM CHIR99021 and 1 uM PD0325901.

### Virus production

Virus were packed in 293T cells. Transfection was performed using Polyethylenimine, and the plasmids pCMV-VSVG (a gift from Bob Weinberg; Addgene plasmid # 8454; http://n2t.net/addgene:8454; RRID:Addgene_8454) (Stewart et al. 2003) and psPAX2 (a gift from Didier Trono; Addgene plasmid # 12260; http://n2t.net/addgene:12260; RRID:Addgene_12260). The viruses were harvested at 48 and 72h post transfection. For the CAS9 virus, the lentiCas9-Blast was used (a gift from Feng Zhang; Addgene plasmid # 52962; http://n2t.net/addgene52962; RRID:Addgene_52962) (Sanjana et al. 2014), and for the knockdown library the Brie pooled library was used (a gift from David Root and John Doench; Addgene #73633) (Doench et al. 2016).

### Generation of CAS9 expressing cells

mESCs were transfected with a lentivirus containing the CAS9 expression vector at increasing multiplicity of infection (MOI) from 0.5 to 4 in four different wells. The cells were then treated with Blasticidin for 7 days, and the well with the highest number of surviving cells was maintained (MOI 4).

### Screen for essential genes using pooled CRISPR library

The Brie knockout library was used in the screen. The library targets 19,674 genes, with 4 gRNAs targeting each gene. In addition, the library contains 1000 control gRNAs which allows for an estimation of random drift effects on gRNA abundance. CAS9 expressing mESC were transfected with lentiviruses containing the Brie pooled library at MOI ~ 0.4. The cells were treated with puromycin for 7 days. Following the antibiotic selection, the cells were allowed to proliferate for additional 10 days. Cells were passaged at days 8, 11, 15 and 18 post transfection. For each passage a minimum number of 31 million cells was retained for sequencing and additional 31 million cells were re-plated, this allowed for a maintenance of adequate library representation (an average of ~400 cells per gRNA).

### DNA extraction, amplification and sequencing

Genomic DNA was extracted from the cells using Zeymo quick-DNA miniprep kit. The extracted DNA was amplified using Q5^®^ High-Fidelity 2X Master Mix, during the amplification adaptors and barcodes were attached. In addition to DNA samples from the transfected cells, a sample of the plasmid library used for the lentivirus production was also sequenced. To minimize variance originating from amplification biases, each sample was amplified in 3-4 different PCR reactions and the products were pooled. The PCR products were purified using AMPure beads. Samples were sequenced on a NextSeq machine (Illumina). Reads were counted by first locating the CACCG sequence that appears in the vector 5′ in all gRNA inserts. The next 20 bases are the gRNA insert, which were then mapped to a reference file of all possible gRNAs present in the library using bowtie2 (Sanjana et al. 2014).

### Identification of essential genes

For all samples, library sizes were normalized using calcNormFactors function in edgeR, which uses the trimmed mean of the M values method (Sanjana et al. 2014). This normalization corrects for under estimation of gRNAs abundances due to the presence of a few highly represented gRNAs. Following the normalization, the log fold change of each gRNA in each proliferation day was calculated relative to the initial counts in the plasmid library. In order to determine which genes are under significant positive or negative selection we used a simulation-based approach that is based on comparing the fold change of gRNAs targeting each gene to the fold change of randomly selected control gRNAs. This approach allows the detection of negative and positive selection that effect only gene targeting gRNAs in the presence of random drift. For each proliferation day we ranked all gRNAs by their representation fold change relative to the library. For each gene we then calculated the sum of ranks of its gRNA across all days. This score was compared to an empirical null distribution generated by randomly selecting 4 control gRNA and calculating their sum of ranks (10,000 simulation). The *P*-value for each gene was calculated by dividing the number of time its score was lower or equal to the scores obtained by the random simulation of control gRNAs. *P*-values were corrected for multiple testing using Benjamini-Hochberg false discovery rate (FDR) procedure. CRISPR score were defined as the sum of ranks for each gene divided by 10^4^.

### Testing for a possible bias arising from amplified genomic regions

Recent studies reported that CAS9-mediated cleavage of amplified genomic regions could cause a DNA damage response that results in cell death (Ruan et al. 2008; Ihry et al. 2018). Thus, low CRISPR scores of genes located in those regions will not necessarily reflect essentiality. In order to test this in our data, we used a sliding window approach to identify contiguous stretches of low CRISPR scores. To this end, for each chromosome we sorted the genes according to their location and used a sliding window of 40 genes. Windows in which more than 30% of the genes had a CRISPR sore in the bottom 5% were further examined. We detected two such gene clusters on chromosome 13 (Figure S7A). When examining the clusters closely, we found that they contain mostly histone genes and that non-histone genes in those regions did not have low CRISPR scores (Figure S7B). Therefore, we concluded that the clusters probably reflect true essentials and not a DNA damage response.

### Overlap of essential genes with previous published dataset

The overlap between genes found to be under significant negative selection (FDR corrected *P* < 0.05) in the screens was tested using Fisher’s exact test. Welch’s t-test was used to test the significance for the difference in the quantitative fold change (day 15 post transfection) between genes found to be under significant or non-significant negative selection in previous screen (Tzelepis et al. 2016). A Combined *P*-value for the two screens were obtained using the sum z (Stouffer’s) method (Stouffer et al. 1949; WHITLOCK 2005). In this method p-values are converted to Z values, combined, and then converted back into *P*-values.

### GO terms and KEGG pathway enrichment analysis

GO terms enrichment of essential genes was performed using the Gorilla tool (Eden et al. 2009) on the background of all genes tested in the screens. GO terms significant at FDR corrected value *P* < 0.05 were summarized using reviGO (Supek et al. 2011). Terms that were not specific and contained more than 20% of the tested genes were removed. In addition, terms for which all the essential genes overlapped in other more enriched terms were removed. Comparison of GO term enrichment between essential genes in the fast and gradual declining group and between genes essential specifically in mESCs or specifically in hESCs relative to genes essential in both mESCs and hESCs was performed using Fisher’s exact test. KEGG pathways enrichment analysis of essential genes was done using the clusterProfiler R package (Yu et al. 2012). Comparison of KEGG pathways enrichment between essential genes in the fast and gradual declining groups was performed using Fisher’s exact test. Comparison of KEGG pathways enrichment between all genes essential to human or mouse ESCs and human phenotypes associated with essential genes was performed using proportion test.

### Analysis of paralog genes

Paralog genes were identified using ensemble biomart (Smedley et al. 2015) and TreeFam (Ruan et al. 2008) databases. The significance of the difference between the CRISPR score distribution of genes with and without a paralog in the top KEGG pathways was tested using a Mann-Whitney test. The enrichment for genes without a paralog for genes in the fast and gradual declining clusters and for non-essential genes was tested using a fisher’s exact test.

### Cluster identification based on gRNA kinetics

Clustering of essential genes in mESC was performed based on the correlation between the depletion rates for all essential genes. The correlation matrix was then used for hierarchical clustering using R hclust function with the default settings. The dendrogram branches were cut to obtain two main clusters.

### Gene expression in ESCs

Gene expression for mESCs was obtained from the study of Tesar et al. (Tesar et al. 2007) and for hESCs from the study of de Leemput et al. (van de Leemput et al. 2014). For mESCs microarray data was normalized using quantile normalization and the mean expression across 3 samples for each gene. For hESCs normalized counts from RNA-seq data were used, and for each gene the mean expression across 4 samples was calculated. To test for significance differences in gene expression between groups we used the Welch’s t-test in case of two groups, and Tukey test for 3 groups.

### Difference in mean half-life between genes in the gradual and fast declining group

Data on protein half-life was obtained from Mathieson et al. (Mathieson et al. 2018) and Schwanhausser et al. (Schwanhäusser et al. 2011). The difference in the mean log10 half-life for genes in the fast and gradual declining groups was determined using Welch’s t-test.

### Comparison of essential genes between mESCs, hESCs and cancer cell lines

Data on essential genes in haploid hESCs was obtained from Yilmaz et al. (Yilmaz et al. 2016); data on essential genes in human cancer cell lines was obtained from Yusa et al. (Tzelepis et al. 2016) and Wang et al. (Wang et al. 2017). Genes essential only in ESCs were defined as genes essential in mESCs and hESCs (FDR corrected *P* < 0.05) but not essential in more than one of the 11 human cancer cell lines. Similarly, genes essential specifically in mESCs were defined as genes not essential in hESCs and not essential in more than one of the 11 human cancer cell lines. Genes essential specifically in hESCs were defined as genes essential in hESCs (FDR corrected *P* < 0.05) not essential in mESCs and not essential in more than one of the 11 human cancer cell lines. The agreement between essential genes in mESCs and the human cells lines was tested by ranking the genes in each cell type by their negative selection level and comparing the rank distribution between genes essential and non-essential in mESCs. The significant differences in the rank distribution was tested using a Mann-Whitney test.

### Differential expression analysis between mESCs and EpiSCs

Microarray gene expression data for 3 mESCs samples and 3 EpiSCs samples was obtained from Zhou et al. (Zhou et al. 2012). The data was normalized by quintile normalization and differential expression was performed using limma (Ritchie et al. 2015). Association with genes significantly upregulated in mESCs relative to EpiSCs was determined using Fisher’s exact test.

### Overlap with mouse embryonic lethal genes and human LoF intolerant genes

A list of genes which have a knockout mouse with a phenotype of prenatal lethality (MP:0002080), abnormal survival (MP:0010769, excluding extended life span), decreased prenatal and postnatal growth (MP:0002089, MP0004196, MP0001731, excluding increased body size, weight gain and diet related phenotypes) was obtained from the mouse Genome Informatics (MGI) database (Bult et al. 2018). The overlap between genes identified as essential in mESC and the list of genes associated with growth and lethality in mice was tested only for genes that have a knockout mouse information in the MGI database. The significance of the overlap was tested using Fisher’s exact test. A list of genes defined as human LoF intolerant was obtained from Lek et al. (Lek et al. 2016). The significance of the overlap between human LoF intolerant genes and essential genes in mESCs and hESCs was calculated using Fisher’s exact test.

### Association of essential genes with human phenotypes

Analysis of human phenotypes for genes essential in mESCs or hESCs was based on phenotypes in the Human Phenotype Ontology (Köhler et al. 2019). The significance of the association was calculated by Fisher’s exact test and *P*-values were corrected for multiple testing by FDR procedure. Significant phenotypes with more than 90% overlap of genes with a more significant phenotype were filtered out. Association with developmental and neurodevelopmental phenotypes for genes essential in mESCs or hESCs, was based on the DDG2P dataset from the deciphering developmental disorders project (Wright et al. 2015). Neurodevelopmental phenotypes were defined as any developmental phenotype involving the brain. The significance of the association was calculated using Fisher’s exact test.

### Gene expression during in vitro human and mouse corticogenesis

Neurodevelopmental disorders risk genes, as defined by the deciphering developmental disorders project (Wright et al. 2015) or genes present in the developmental brain disorders database (tires 1 & 2) (Gonzalez-Mantilla et al. 2016), were clustered according to their expression patterns during *in vitro* corticogenesis of mouse (Hubbard et al. 2013) or human (van de Leemput et al. 2014) cells. Clustering was performed using R hclust function with the default settings (the complete method).

## Acknowledgments

We thank Nissim Benvenisty, Alana Amelan, and Eran Meshorer for valuable comments on the manuscript. This research was supported by the Israel Science Foundation (grant no. 575/17) and by the Israel Science Foundation Broad institute Joint Program (grant no. 2612/18).

